# Predicting Parkinson’s disease progression using MRI-based white matter radiomic biomarker and machine learning: a reproducibility and replicability study

**DOI:** 10.1101/2023.05.05.539590

**Authors:** Mohanad Arafe, Nikhil Bhagwat, Yohan Chatelain, Mathieu Dugré, Andrzej Sokołowski, Michelle Wang, Yiming Xiao, Madeleine Sharp, Jean-Baptiste Poline, Tristan Glatard

## Abstract

**Background:** The availability of reliable biomarkers of Parkinson’s disease (PD) progression is critical to the understanding of the disease and development of treatment options. Magnetic Resonance Imaging (MRI) provides a promising source of PD biomarkers, however, neuroimaging results have been shown to be markedly sensitive to analytical conditions and population sampling, which motivates investigations of their robustness. This study is part of a project to investigate the replicability of 11 structural MRI measures of PD identified in a recent review.

**Objective:** This paper attempts to reproduce (similar data, similar analysis) and replicate (variations in data and analysis) the design of the machine learning (ML) model described in [1] to predict PD progression from T1-weighted MRIs.

**Methods:** We used the Parkinson’s Progression Markers Initiative dataset (PPMI, ppmi-info.org) used in [1] and we followed as closely as possible the original methods. We also investigated slight methodological variations in cohort selection, feature extraction, ML model design, and evaluation techniques.

**Results:** The Area under the ROC Curve (AUC) achieved by our model closely reproducing the original study remained lower than 0.5. Across all tested models, we obtained a peak AUC of 0.685, which is better than chance performance but remained lower than the AUC value of 0.795 reported in [1].

**Conclusion:** We managed to train a model that predicts disease progression with a performance better than chance on a cohort extracted from the PPMI dataset, using methods adapted from [1]. However, the performance of this model remains substantially lower than the one reported in [1]. Our difficulties to reproduce or replicate the original work are likely explained by the relatively low sample size in the original study. We provide recommendations on how to improve the reproducibility of MRI-based ML models of PD in the future.

## Introduction

Identifying biomarkers that can predict the progression of Parkinson’s disease (PD) is essential to support the development of new therapies and track responses to these therapies. T1-weighted magnetic resonance imaging (MRI) is a promising source of PD biomarkers as it non-invasively provides detailed information about disease-related changes to brain structure. Among MRI analysis methods, machine learning (ML) techniques are growing in popularity due to their excellent predictive ability. For example, a recent review [2] identified 110 studies that used ML for PD prediction, among which 79 used some form of brain imaging data including MRI. However, despite their potential, MRI-based measurements of PD have yet to be widely adopted in clinical and research settings, in part due to the lack of reliability, robustness, and reproducibility of such measures.

As with many other disciplines, the reproducibility and replicability of neuroimaging findings has been under increasing scrutiny. For instance, the study in [3] asked 70 independent teams to test nine different hypotheses using the same functional MRI dataset and highlighted substantial discrepancies among the teams for five of these hypotheses. Moreover, the work in [4] showed that longitudinal MRI-based measures of cortical thickness led to conflicting results for PD progression. Further, studies have shown that measurements of anatomical volume and cortical thickness are affected by the software versions, workstation types, and versions of operating system used [5]. ML itself is also subject to reproducibility concerns. For instance, the study in [6] attempted to reproduce 20 ML-based studies from 17 fields and revealed several pitfalls related to data leakage between the training and test sets, resulting in a substantial number of failed replications. Therefore, the reproducibility and replicability of ML-based MRI studies need to be investigated.

The terms reproducibility and replicability are distinct and may cause confusion. In this article, reproducibility refers to the ability to obtain the same results as in the original study by running the same software on the same input data, whereas replicability refers to the ability to obtain results comparable to the original ones by repeating the experiment with different data and software [7]. In practice, the distinction between reproducibility and replicability is blurry, and these terms have to be understood as the extrema of a continuous range of variations from the original study.

This paper is an attempt to both reproduce and replicate the study in [1]. Among MRI-based studies of PD that used ML (e.g., [8] [9]), the work of [1] is particularly interesting as it relies on an openly accessible dataset — the Parkinson’s Progression Markers Initiative (PPMI [10]) — and it targets the clinically-relevant problem of predicting PD progression. In [1], the authors trained a support vector machine (SVM [11]) classifier using whole-brain white matter (WM) and clinical features to predict the progression of PD over 3 years. To do so, they segmented WM masks from T1-weighted MRI scans of n=144 patients (72 stable, 72 progressive), extracted radiomics features from the WM, trained a classifier with these features, and evaluated its capability to predict PD progression measured by the Hoehn and Yahr Scale (HYS), resulting in a AUC of 0.795. The work in [1] also developed a joint model that combined radiomics features with clinical features, and achieved a slightly better performance with an AUC of 0.836. Our study focuses primarily on imaging biomarkers, hence, we focused on evaluating the reproducibility and replicability of the radiomics model only.

## Methods

The authors of [1] trained a linear SVM from radiomics features to predict disease progression measured with the Hoehn and Yahr Scale (HYS). The resulting model achieved an AUC of 0.795 and a relative standard deviation (RSD) of 3.23 % across 100 bootstrap samples. Our objective was to assess the reproducibility and replicability of the original study. For the reproducibility experiment, we attempted to reproduce the cohort, feature set, model, and evaluation technique using the methods described in [1]. We then compared the AUC and RSD of our reproducibility experiment to the results reported in [1]. For the replicability experiment, we created several variations of the methods described in [1] by creating multiple cohorts from the PPMI dataset, various feature sets, and two different evaluation techniques. We tested each possible combination of these variations and compared the resulting AUC values to those achieved in [1]. Figure 1 shows a summary of all the configurations tested.

**Fig. 1.**
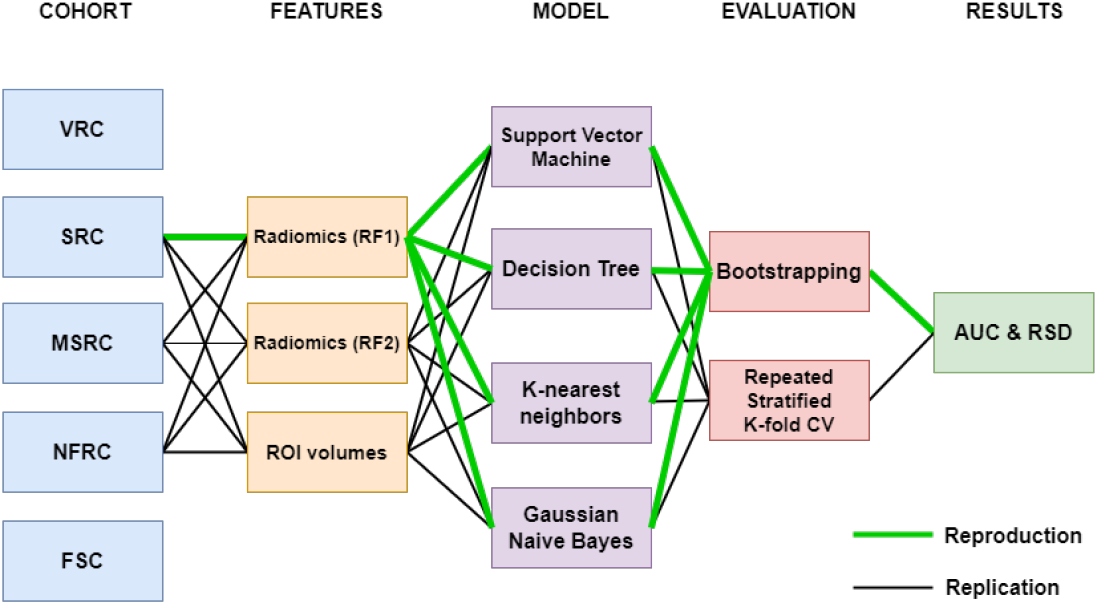
Configurations tested in the reproduction and replication experiments. Cohorts include the Verio Reproduction Cohort (VRC), Siemens Replication Cohort (SRC), Multiple Scanner Replication Cohort (MSRC), No Filter Replication Cohort (NFRC) and the Functional State Cohort (FSC). Features include the mapped radiomic features (RF1), selected radiomic features (RF2), and volumes of regions of interest (ROI volumes). The green line represents the configuration for the reproducibility experiment while the black lines represent the different configurations of the replicability experiment.

### Cohort construction

The original study in [1] included PD patients matched by age, sex, and baseline HYS value across each group from the PPMI dataset. MRI data was collected from 32 international sites using a Siemens Verio 3T MRI machine. The protocol used for data acquisition was standardized by PPMI protocols and included the following parameters: repetition time = 2300 ms, echo time = 2.98 ms, inversion time = 900 ms, slice thickness = 1 mm, field of view = 256 mm, and matrix size = 240 × 256. Each patient was evaluated through a baseline visit that included MRI acquisition and clinical examination, and a follow-up clinical examination after 3 years (36 months). Patients with HYS values higher at follow-up than at baseline were included in the progressive set (n=72) and patients with the same HYS value at follow-up and at baseline were included in the stable set (n=72). Data was collected after approval of the local ethics committees of the PPMI’s participating sites. All participants provided written informed consent.

We constructed a total of 5 cohorts by applying different combinations of filters to the PPMI database (Table 1). For all cohorts, we filtered the PPMI database for patients meeting the following inclusion criteria:

- **C1**: received a diagnosis of idiopathic PD;
- **C2**: has a pair of visits spaced 3 years apart, with a T1-weighted MRI available at the first visit;
- **C3**: has HYS values available at both visits.

**Table 1.** Summary of PPMI filters used in the 5 constructed cohorts.

|  | VRC | SRC | MSRC | NFRC | FSC |
| --- | --- | --- | --- | --- | --- |
| Research Group | PD | PD | PD | PD | PD |
| Acquisition Type | 3D | 3D | 3D | 3D | 3D |
| Field Strength | 3T | 3T | 3T | any | 3T |
| Slice Thickness | 1mm | 1mm | 1mm | $1\text{mm} \leq 1.2\text{mm}$ | 1mm |
| Manufacturer | Siemens | Siemens | any | any | Siemens |
| Manufacturer model | Verio | any | any | any | any |
| Weighting | T1 | T1 | T1 | T1 | T1 |

Our study was conducted in accordance with the Declaration of Helsinki and was exempt from the Concordia University’s Research Ethics Unit. We did not have access to any personal information such as patients names or addresses. By using the PPMI dataset we accepted its Data Usage Agreement which in particular prevents us from redistributing the data or sharing patients identifiers publicly.

Data was collected by PPMI between 2010 and 2023, we have accessed the database November 22^nd^, 2022. Since PPMI is a longitudinal dataset, a single patient could potentially meet the criteria for both groups at different points in the study, or be included multiple times in the same group. Therefore, after constructing the cohort, we also verified that (i) both groups (stable and progressive) were of equal size, (ii) no patient was present more than once in each group, and (iii) no patient was present in more than one group. We generated two groups of cohorts which will be referred to as the <u>reproduction cohort</u> and the <u>replication cohorts</u>.

#### Reproduction cohort

In our reproducibility experiment, our objective was to build a cohort as close as possible to the one in [1]. The resulting cohort will be referred as the Verio Reproduction Cohort (VRC) as it is the only cohort built with the manufacturer model set to Verio (Table 1). We built the VRC using all the filters that we could extract from the methods section of [1].

#### Replication cohorts

To evaluate the sensitivity of the predictions to data variations, we constructed 3 cohorts with increasingly permissive filters. The first cohort included patients scanned using a Siemens manufactured MRI machine but not necessarily with the Verio model. We will refer to this cohort as the Siemens Replication Cohort (SRC). Compared to the VRC, the SRC has a slightly more permissive manufacturer model filter, which is meant to accommodate variations in manufacturer model descriptions throughout the PPMI study. We constructed two more cohorts with increasingly permissive filters. The Multiple Scanner Replication Cohort (MSRC) includes patients scanned with any scanner manufacturer, and the No Filter Replication Cohort (NFRC) includes patients scanned with any field strength and a slice thickness between 1 mm and 1.2 mm.

We also constructed a functional state cohort (FSC) that takes into account the functional state of patients at each visit. The PPMI protocol requires that clinical assessments be conducted twice per visit in different functional states (“ON state” vs “OFF state”). The methods in [1] did not mention if patients in their cohort were in the ON or OFF state. The variables related to a patient’s functional state during a visit are reported in MDS-UPDRS Part III evaluations and include:

- PDSTATE (ON/OFF): the current functional state of the patient
- PDTRTMNT (0/1): 1 if the participant is on PD medication or receives deep brain stimulation, 0 otherwise
- PDMEDTM: time of most recent PD medication dose
- PDMEDDT: date of most recent PD medication dose

In the FSC, we modified inclusion criterion **C3** so that HYS measures of a given patient were obtained with the same PDSTATE (ON or OFF) at both visits. This was meant to ensure that HYS measures were consistently obtained between visits and were therefore comparable. Moreover, the MDS-UPDRS Part III evaluations available in PPMI contain inconsistencies and missing data that we corrected as described in https://github.com/LivingPark-MRI/livingpark-utils/blob/main/livingpark_utils/notebooks/pd_status.ipynb.

For each sub-cohort, we used the list of patients returned by the PPMI query and kept those that have a pair of MRI visits spaced 3 years apart. Furthermore, we matched patients from both groups based on age, sex and baseline HYS value.

### MRI Feature extraction

We extracted two sets of image features for each cohort. The first set of features (**RF1** and **RF2**) was radiomics-based as per the methods reported in [1]. The second set of features (**ROI**) consisted of WM, gray matter (GM) and ventricle volumes measured from known ROIs involved in Parkinsonian syndromes [12].

#### Segmentation of T1-weighted images

For **RF1** and **RF2**, we used the Segmentation module of Statistical Parametric Mapping (SPM; https://www.fil.ion.ucl.ac.uk/spm/software/spm12 [13]) version 12 that was also the segmentation method used in [1]. We used SPM12’s default parameters to get the tissue probability masks and build a WM binary mask for each patient. For **ROI**, we used FreeSurfer v6.0 (recon-all tool with parameter -brainstem) to extract the ROI volumes needed, as done in [12].

#### Quality control

In [1], two experienced neuro-radiologists used ITK-snap to manually edit WM segmentations. The modifications included (i) removal of non-brain tissue, brain stem and cerebellum and (ii) correcting segmentation errors in WM tissues. We used 3D Slicer v.5.0.3 to visualize and assess the quality of WM segmentations produced by SPM12. For each MRI scan, we reviewed the axial, coronal and sagittal slices. Data were excluded if it met at least one of the following criteria:

- There is WM outside of the segmented WM mask;
- There is GM inside the segmented WM mask;
- The MRI has any common artifacts;
- The MRI has a low signal-to-noise (SNR) ratio.

We processed and applied these exclusion criteria to all the images meeting inclusion criteria **C1, C2** and **C3**, leading to a total of 7 excluded images. In addition, 11 images could not be processed with FreeSurfer due to execution errors. We applied the cohort selection filters to the resulting set of images.

#### Radiomic features

The A.K. software (Artificial-Intelligent Radio-Genomics Kits; GE Healthcare, Chicago, IL, USA) used in [1] is not publicly available. Therefore, we used PyRadiomics [14], an open-source Python package for the extraction of radiomics features. PyRadiomics can extract a total of 56 features relevant to our study, including 24 gray level co-occurrence matrix (GLCM) features, 16 gray level size zone matrix (GLSZM) features and 16 gray level run length matrix (GLRLM).

In [1], the authors extracted a total of 378 features, including 42 histograms features, 10 Haralick features, 9 FormFactor features, 126 GLCM features, 180 GLRLM features, and 11 gray level region matrix features (GLZSM). From these 378 features, the authors used the maximum relevance minimum redundancy (mRMR) algorithm to extract the following top 7 features and train the model:

- Feature 1: GLCMEntropy AllDirection offset1
- Feature 2: RunLengthNonuniformity angle45 offset7
- Feature 3: Correlation angle45 offset1
- Feature 4: HaralickCorrelation angle90 offset4
- Feature 5: ShortRunEmphasis angle0 offset7
- Feature 6: HaralickCorrelation AllDirection offset7
- Feature 7: Inertia AllDirection offset4

We extracted two sets of features using PyRadiomics. The first set, **RF1**, included 5 PyRadiomics features that best match the 7 A.K software features from [1], namely:

- Feature 1: Joint Entropy
- Feature 2: Run Length Non Uniformity
- Feature 3 / Feature 4 / Feature 6: Correlation
- Feature 5: Short Run Low Gray Level Emphasis
- Feature 7: Contrast

The mapping between A.K software and PyRadiomics features is not exact. Indeed, the A.K software, unlike PyRadiomics, provides every feature at a specific angle and offset. In PyRadiomics, for each feature class, the value of a feature is calculated for each angle separately, after which the mean of these values is returned. The exact definitions of these features are available in the PyRadiomics documentation (https://pyradiomics.readthedocs.io/en/latest/features.html) and in the supplementary material of [1], Table S2.

The second set of radiomics features, **RF2**, included the top 7 features from the 56 ones extracted by PyRadiomics. We selected the top 7 features using the mRMRe R package [15] also used in [1], with R v4.2.1 and k=7.

#### Volumes of regions of interest

We used FreeSurfer v6.0 to extract 13 ROIs that contribute to Parkinsonian syndromes as shown in [12]. Those include the midbrain, pons, putamen, posterior putamen, caudate, thalamus, pallidum, precentral cortex and insular cortex in the gray matter, the superior cerebellar peduncle, and the cerebellum white matter including the middle cerebellar peduncles in the white matter and the third and fourth ventricles. Every region’s volume was used as input features in the ML models.

#### Feature normalization

We normalized and centered the features using scikit-learn’s StandardScaler, resulting in the following transformation:

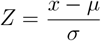

where *μ* is the mean of the training samples and *σ* is the feature standard deviation in the training set. We then applied this transformation to the test set, reusing the mean and standard deviation values learned from the training set.

### Prediction model

To predict disease progression, the authors of [1] trained a linear SVM based on the 7 top features extracted and selected from segmented WM masks of PD patients. The authors compared the SVM with three other machine learning methods, including Gaussian Naive Bayes (GNB), k-nearest neighbours (KNN), and decision tree (DT) classifiers. Since the methods of [1] did not mention the name and values of the classification hyper-parameters that were optimized, we optimized the usual parameters for these classifiers using the ranges in Table 2. We implemented the models using scikit-learn v1.1.3 and Python v3.10.4.

**Table 2.**
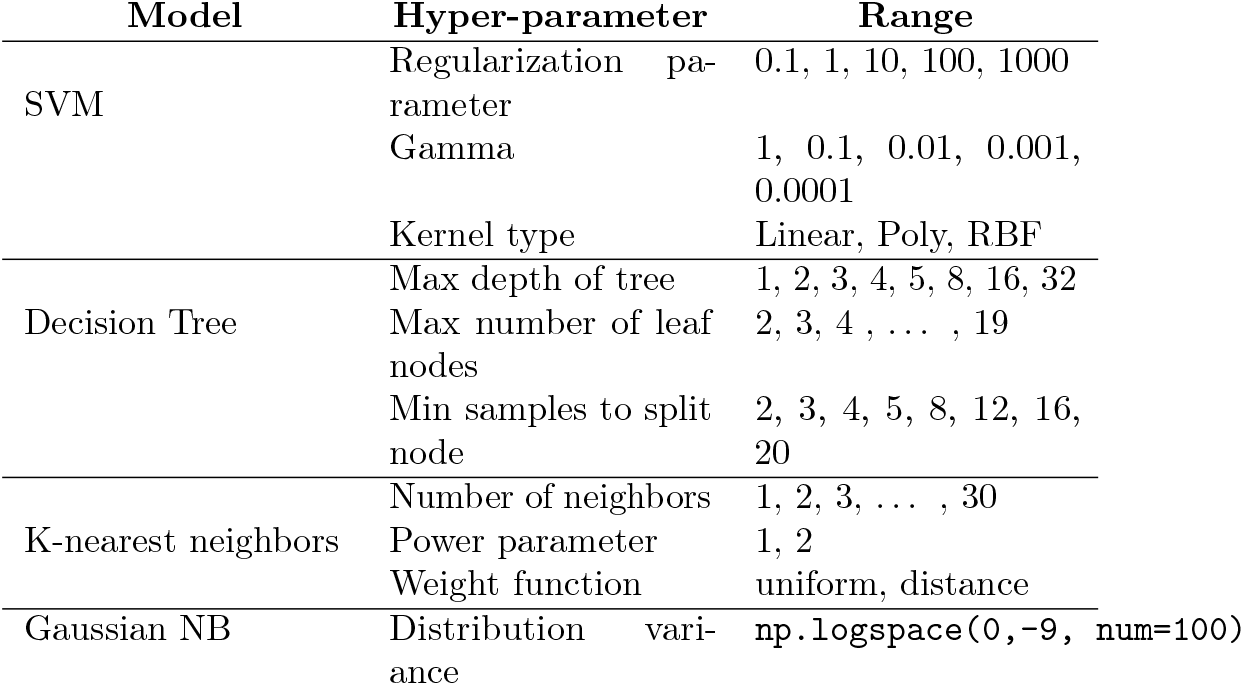
Ranges used in hyper-parameter optimization.

#### Model selection and evaluation using bootstrap

In [1], the authors used a bootstrap approach to compare classifiers and optimize their hyper-parameters. To reproduce this process, we split the dataset into training (100 patients) and test (44 patients) sets having matched the HYS values of the patients in training and test sets. The size of the training and test sets matched the ones reported in [1]. We implemented model selection using 100 iterations of a bootstrap sampling loop applied to the training set. Each iteration randomly selected (with replacement) 50 patients, normalized the features for these 50 patients, fitted the models to these 50 patients, and measured the AUC of the models on the remaining patients. To measure the stability of the models across the 100 bootstrap samples, we computed the RSD defined as:

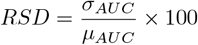

where *σ*_*AUC*_ and *μ*_*AUC*_ are the standard deviation and mean of the AUC values obtained on the 100 bootstrap samples. We selected the model with the lowest RSD and applied it to the test set.

#### Model selection and evaluation using cross-validation

As part of the replication experiment, we also implemented a Stratified K-fold cross-validation (CV) loop similar to the one in [12] and more commonly seen in the ML literature than the bootstrap loop mentioned previously. We first split the cohort into training (100 patients) and test (44 patients) sets randomly. For model selection, we applied to the training set a CV loop including 50 repetitions of a 5-fold CV stratified with the target variable (progressive/stable class) to preserve its distribution across folds. For each fold, we normalized the features using the standard scaler mentioned above, selected hyper-parameters based on the performance of the validation set and reported the AUC computed on the test set using the model that produced the best average AUC in the validation fold. We implemented this CV loop independently for the SVM, GNB, kNN, and DT, with a scikit-learn validation pipeline, using scikit-learn’s RepeatedStratifiedKFold function with 5 splits and 50 repetitions, and scikit-learn’s GridSearchCV function with the parameters in Table 2.

### Infrastructure & code availability

We used Pandas v.1.4.3 and Numpy v1.22.4 to construct the cohorts. The extraction of WM using SPM12 was carried out by running Boutiques tool with DOI 10.5281/zenodo.6881412 using Docker v20.10.12 and Boutiques v0.5.25 [16]. The construction of the cohort, extraction of radiomics features, and training of the ML models were performed on a local computer using Ubuntu OS version 22.04. The FreeSurfer segmentations, on the other hand, were executed remotely through the Cedar cluster of the Digital Research Alliance of Canada, which operated on CentOS Linux 7 (Core) operating system with Linux kernel v3.10.0.

All our methods are available in a publicly available notebook (https://github.com/LivingPark-MRI/shu-etal). To comply with PPMI’s Data Usage Agreements that prevent users to re-publish data, the notebook queries and downloads data directly from PPMI. Since PPMI does not have a data access API, we developed our own Python interface to PPMI using Selenium, a widely-supported Python library to automate web browser navigation. Using this interface, the notebook downloads PPMI study and imaging files to build the cohorts and train the ML models. The utility functions to download and manipulate PPMI data are implemented in LivingPark utils, a Python package available on GitHub (https://github.com/LivingPark-MRI/livingpark-utils).

## Results

### Cohorts

Table 3 summarizes the demographics of the reproduction and replication cohorts that we obtained. Although we built the VRC using the same PPMI filters as in [1], we were not able to reproduce the original cohort due to a shortage of subjects scanned with a Verio scanner. In fact, when we performed the query, we found a total of 29 visit pairs for progressive patients with HYS=1, 98 visit pairs for progressive patients with HYS=2, 0 visit pairs for stable patients with HYS=1, and 66 visit pairs for progressive patients with HYS=2. Using these visit pairs, we were not able to match the number of patients in [1] while ensuring that a given patient appears in at most one group.

**Table 3.**
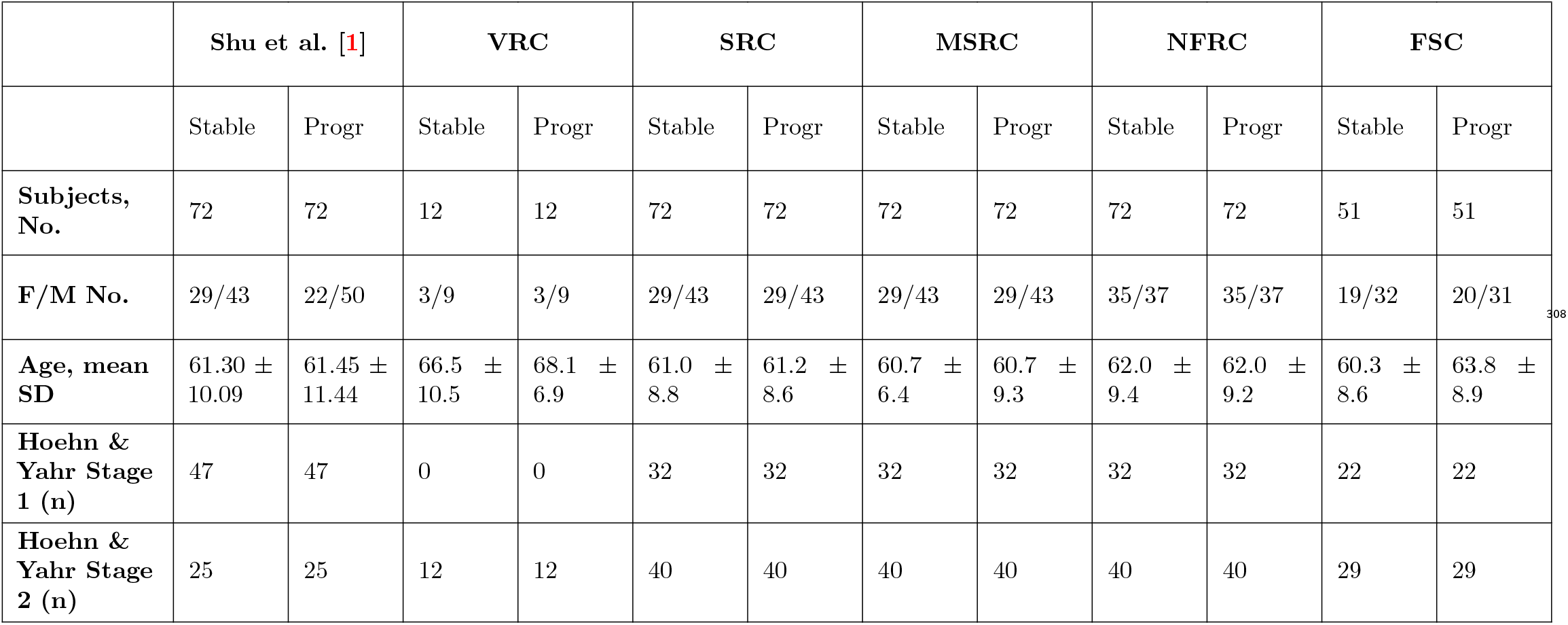
Summary of the reproduction and replication cohorts constructed.

The SRC is the closest cohort to the one in [1] that we were able to build. As in [1], the SRC includes 72 progressive and 72 stable patients scanned with a Siemens manufactured MRI machine. However, the patient breakdown by HYS value differs from [1] in each group: in the SRC, both groups have 32 patients with baseline HYS=1 and 40 patients with baseline HYS=2 whereas in [1] these numbers are respectively 47 and 25. The age and F/M balance in the SRC are comparable to the one reported in [1], with 29 females and 43 males per group, an average age of the stable group of 61.0±8.8, and an average age of the progressive group of 61.1±8.6. Finally, it should be noted that out of 144 patients in the SRC, 96 have a different value of PDSTATE (ON/OFF) at their baseline and follow-up visits.

The MSRC includes all the patients meeting the SRC’s inclusion criteria except for the MRI machine used. In the MSRC, 132 patients have been scanned with a Siemens machine, 4 with a GE Medical Systems machine, 8 with a Philips machine and 1 with an unknown machine. There are 29 females and 43 males per group. The average age of the stable group is 60.7 yrs ± 9.4 and the average age of the progressive group is 60.7 yrs ± 9.3. Both groups have 40 patients with baseline HYS=1 and 32 patients with baseline HYS=2. Finally, 103 patients have a different value of PDSTATE (ON/OFF) at their baseline and follow-up visits.

The NFRC included patients with an MRI of any field strength and slice thickness between 1 mm and 1.2 mm. In the NFRC, 108 patients have been scanned with a Siemens machine, 19 with a GE Medical Systems machine, 15 with a Philips machine and 2 with unknown scanners. There are 35 females and 37 males per group. The average age of the stable group is 62.0 ± 9.4 and the average age of the progressive group is 62.0 ± 9.2. Both groups have 40 patients with baseline HYS=1 and 32 patients with baseline HYS=2. Finally, 100 patients out of the 144 have a different value of PDSTATE (ON/OFF) at their baseline and follow-up visits. The FSC includes an additional filter to only keep visit pairs with consistent values of PDSTATE (ON/OFF) between the baseline and follow-up visits. The FSC only includes 102 patients and therefore does not reproduce the sample size in the cohort of [1]. In total, we could only find 22 patients with HYS=1 and 29 patients with HYS=2 in each group, totalling 102 patients in the cohort.

### Extracted features

The second set of features, **RF2**, consisted of the top 7 features selected by the mRMR algorithm applied to the 56 features available in PyRadiomics. The top 7 features identified varied by cohort and are reported in Table 4. The cluster shade and Size-Zone Non-Uniformity Normalized appear in all three cohorts. Notably, none of the features extracted in [1] appear in any of the cohorts.

**Table 4.**
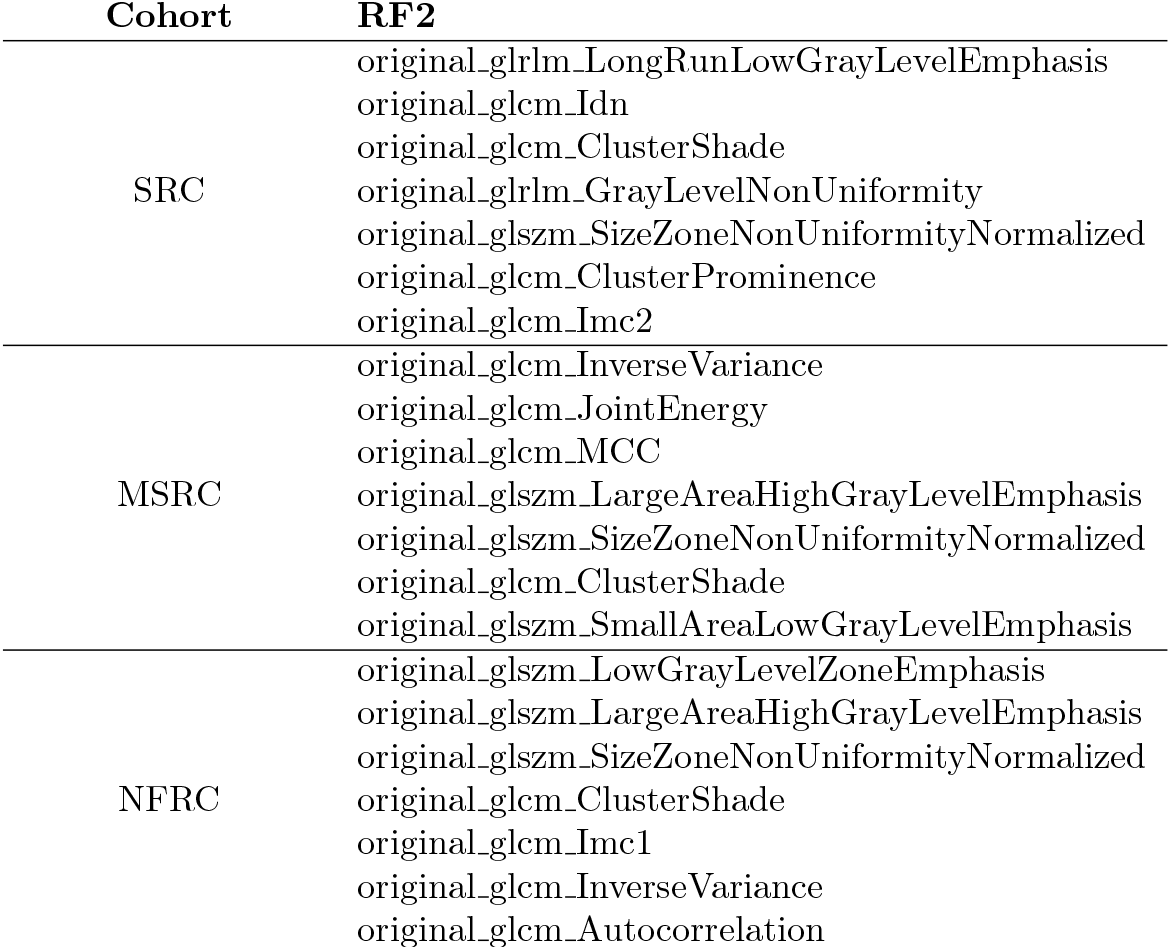
Feature extraction (RF2) per cohort using mRMRe (k=7)

### Reproducibility experiment

Our first objective was to reproduce the pipeline of [1]. For this reproducibility experiment, we used the SRC since it is the closest cohort to the one in [1] that we could create. We used the 5 radiomic features (**RF1**) extracted with PyRadiomics (Figure 2). We trained the 4 models mentioned previously (SVM, kNN, GNB, DT), optimized the hyperparameters as described in Table 2, and used the bootstrap evaluation approach.

**Fig. 2.**
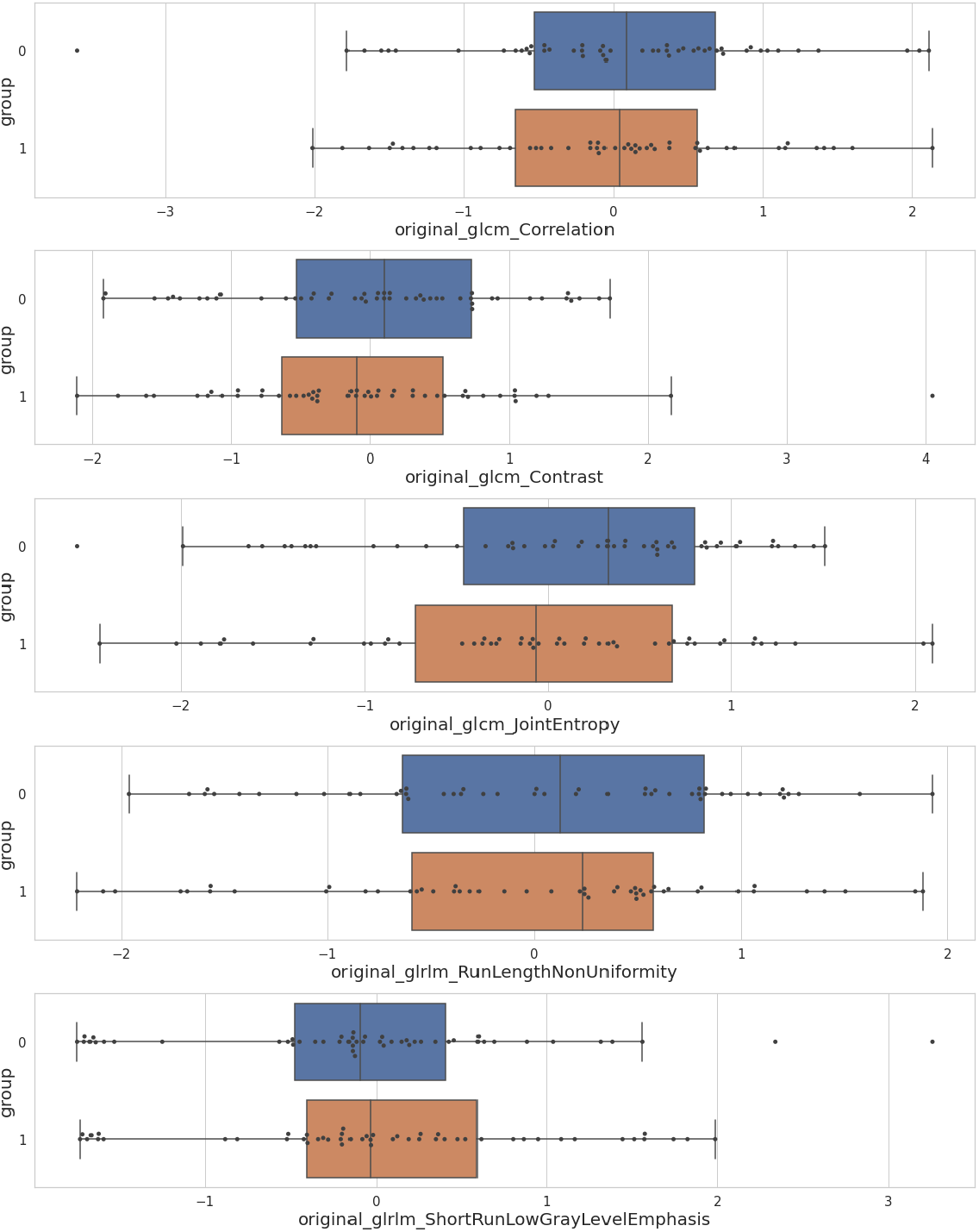
Distribution of **RF1** features in the training set in the SRC. 0: stable group; 1: progressive group.

The evaluation results for all the models on the validation set are reported in Table 5. After hyper-parameter tuning, none of the models achieved an average AUC higher than 0.501 on the validation set. The Decision Tree model had the lowest RSD and was therefore selected for evaluation on the test set (with hyperparameters max depth=1, max leaf nodes=2, min samples split=2) on which it achieved an AUC of 0.523, only slightly better than chance performance. We were not able to reproduce the AUC of 0.795 reported in [1].

**Table 5.** AUC and RSD values of each model’s best performer (defined as model with lowest RSD) in the validation set. The Decision Tree achieved the lowest RSD and was therefore selected for the test set in which it achieved an AUC of 0.523.

|  | AUC | RSD (%) |
| --- | --- | --- |
| **SVM** | 0.456 | 10.819 |
| **Decision Tree** | 0.473 | **7.443** |
| **kNN** | 0.501 | 10.182 |
| **Gaussian NB** | 0.441 | 12.387 |

Figure 3 shows the ROC of every model in the validation set. The observed average AUCs reached a maximal value of 0.501. The maximal AUC value observed across all bootstrap samples was 0.657, reached by the SVM model. The difference between the average and maximal AUC values achieved by the models suggests that, if the PyRadiomics features are indeed equivalent to the original ones, the performance values reported in [1] might have been the result of a random sampling artifact whereby the validation procedure picked a test set favorable to the classifier performance.

**Fig. 3.**
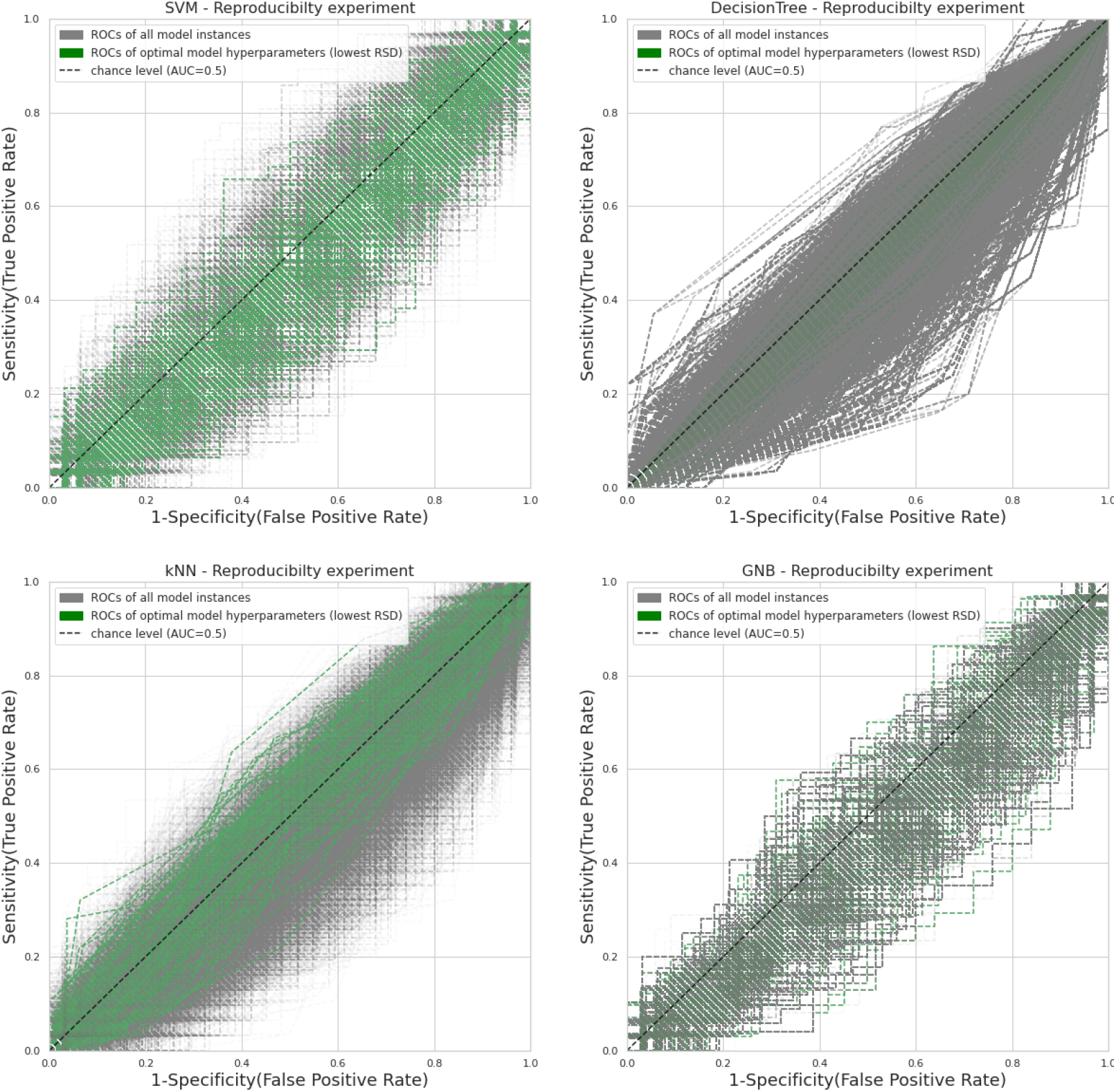
ROC curves per classifier. Gray lines represent ROC curves for model instances over different bootstrap iterations and hyper-parameters. The green curves show the 100 iterations of the hyper-parameter configuration that produced the lowest RSD across bootstrap samples.

### Replicability experiment

Our second objective was to test the replicability of the model described in [1] by using several cohorts and feature sets variations. For every cohort, we trained the four models with the 3 feature sets (**RF1, RF2**, and ROI volumes) using the repeated stratified K-fold CV loop, resulting in 12 sets of results per cohort. The evaluation results for all the models are reported in Table 6 and in Figure 4. While a few configurations reached a mean AUC greater than 0.6, none of them reached the AUC of 0.795 reported in [1]. Most of the results were under chance level (AUC=0.5).

**Table 6.**
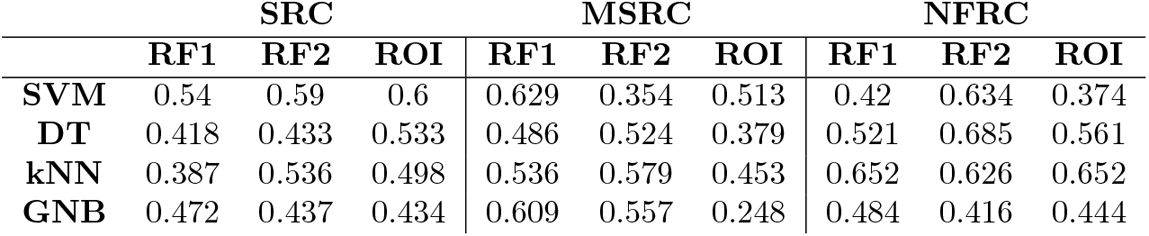
Model AUCs obtained on the test set for the the replicability experiment.

|  | SRC |  |  | MSRC |  |  | NFRC |  |  |
| --- | --- | --- | --- | --- | --- | --- | --- | --- | --- |
|  | RF1 | RF2 | ROI | RF1 | RF2 | ROI | RF1 | RF2 | ROI |
| **SVM** | 0.54 | 0.59 | 0.6 | 0.629 | 0.354 | 0.513 | 0.42 | 0.634 | 0.374 |
| **DT** | 0.418 | 0.433 | 0.533 | 0.486 | 0.524 | 0.379 | 0.521 | 0.685 | 0.561 |
| **kNN** | 0.387 | 0.536 | 0.498 | 0.536 | 0.579 | 0.453 | 0.652 | 0.626 | 0.652 |
| **GNB** | 0.472 | 0.437 | 0.434 | 0.609 | 0.557 | 0.248 | 0.484 | 0.416 | 0.444 |

**Fig. 4.**
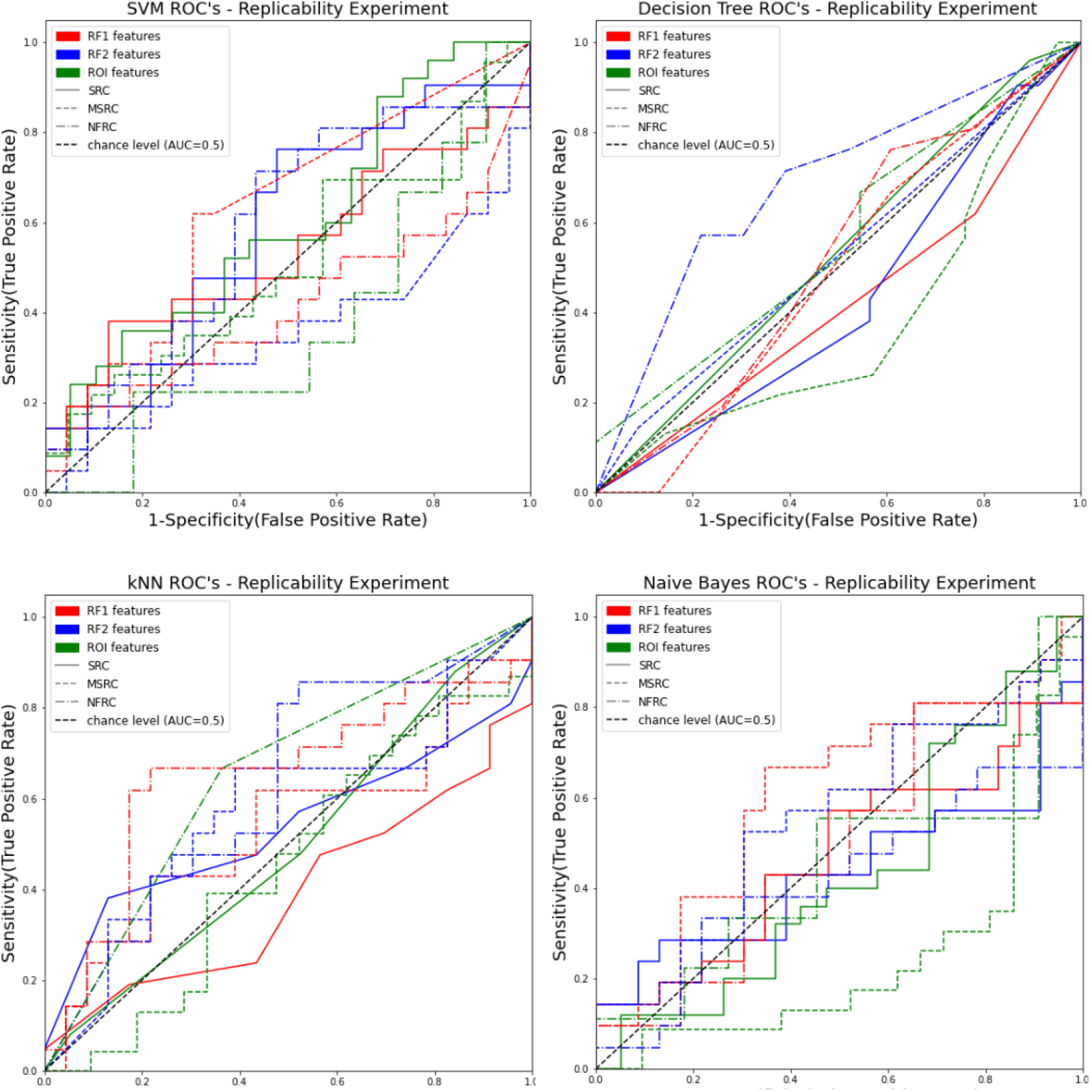
ROC curves of replicability experiment across all cohorts and feature sets on the test set.

## Discussion

We investigated the reproducibility and replicability of the MRI-based ML model of PD progression described in [1]. The performance of the model trained with the methods and cohort closest to the ones described in [1] (reproducibility experiment) did not exceed chance level. When introducing variations in the cohort selection, feature selection, and model evaluation (replicability experiment), several models had AUC values greater than 0.6, with a maximum of 0.685. In summary, we managed to train a model that predicts disease progression with a performance higher than chance on a cohort extracted from the PPMI dataset, using methods adapted from [1]. From that point of view, we could conclude that the replication experiment was successful.

However, the peak performance of our models remained significantly lower than the AUC of 0.795 reported in [1]. We therefore did not replicate nor reproduced the original prediction performance.

While attempting to reproduce and replicate the original study, difficulties occurred in all phases of the analysis, namely cohort construction, feature extraction, and ML model construction. We first attempted to reproduce the exact cohort used in [1] by applying the same filters as described in the original paper, which did not produce enough samples to reconstruct the original cohort. Instead, the Siemens Replication Cohort (SRC) was used as the main reference since it matched the sample size and class distribution in [1]. However, major differences remained between the SRC and the cohort used in [1], in particular regarding the distribution of HYS values (1/2) in each group, which was 32/40 in the SRC and 47/25 in [1]. This disparity in HYS values is a potential reason explaining the observed discrepancies between our models and the ones in [1], as disease progression critically relates to disease stage.

The withdrawal of participants from the PPMI study might be a possible reason why we could not reproduce the cohort of [1]. However, according to the PPMI protocol (section 22), data collected before the withdrawal of a participant should not be removed from the dataset. Another possible explanation could be due to changes in the PPMI user interface or data schema, which may have led to different query results. For instance, during our experiments we found out that approximately 800 images labeled as Proton Density weighted in the image search tool were actually T1-weighted images, which obviously impacted the result of cohort searches. The introduction of the “Mfg Model” (manufacturer model) filter may also have impacted our ability to search the database, as not all patient metadata may have been updated with the Mfg Model information.

While we acknowledge the technical and ethical challenges associated with the public release of multimodal longitudinal data for large, multicentric cohorts, and the permanent need for data curation, the outcome of our reproducibility experiment leads us to urge data-sharing initiatives — including PPMI — to adopt clear data versioning practices and to document data releases. Several mechanisms and tools currently exist to provide these functionalities for neuroimaging data. For instance, DataLad [17] is an extension of the Git version control system to keep track of data file versions, and has been adopted by major data sharing initiatives including OpenNeuro [18] or the Canadian Open Neuroscience Platform [19]. Other initiatives, such as the Human Connectome Project [20], version and periodically release data through their web portals. With clear data releases, we might have identified and possibly resolved the discrepancies between the version of the PPMI dataset that was accessible to us at the time of the experiment, and the version used in [1].

Reproducing the feature extraction pipeline of [1] also raised challenges as we did not have access to the code used in the original study, and the methods section lacked details to fully reproduce the original protocols. In particular, the original study involved a manual correction step of the white-matter masks, which we were unable to replicate. Instead, we quality controlled the white-matter masks through visual inspection, which might have introduced differences with the original study. Moreover, the A.K software used in [1] to extract radiomics features is not publicly accessible and we could not obtain access to it. Instead, we used the PyRadiomics open-source library to extract similar radiomics features, which might have introduced differences with the original study. Overall, the challenges encountered while reproducing the feature extraction step of [1] could have been mitigated by (1) a clear documentation of all manual steps involved in data pre-processing, including quality control and other corrections, (2) the exclusive use of publicly available software.

The ML model trained with the methods reported in [1] yielded under-performing results, achieving AUC values that did not exceed chance level. We believe that this poor performance is due to the small size (n=100) of the training set given the difficulty of the prediction task. Indeed, our models exhibited clear patterns of overfitting that likely resulted from the sparsity of the feature and class distribution samples in the dataset. For instance, the most stable decision tree model (RSD=7.4%) had a maximum depth of 1, which is clearly too shallow to predict disease progression. Deeper decision trees likely overfitted the training set, resulting in larger RSD values still not exceeding chance AUC. The relative small size of the test set (n=44) is also likely to explain the observed fluctuations of classification performance, as illustrated by the jagged ROC curves in Figure 3, as well as the inconsistent performance of classifiers and feature sets across cohorts. In summary, we believe that the differences in classification performance between our models and the one reported in [1] originate in random sampling artifacts resulting from the low sample size. We recommend that future MRI-based ML models of PD prediction be trained on substantially larger samples, to improve the reproducibility of their performance evaluations.

The fact that some of the models included in our replicability experiment performed better than chance on the test set (AUC *≥* 0.6) suggests that the methods described in [1] might produce well-performing classifiers when the feature and class distributions in the training set correctly approximate the ones in the test set. In that sense, our results could be interpreted as a successful replication of the original study. However, given the small size of the test set, it is equally plausible that our reported AUC values are also the result of a random fluke. Here again, further experimentation with larger samples is required.

The challenges faced when attempting to reproduce the original ML model might also come from differences in ML validation pipelines. In particular, the bootstrap validation approach used in [1] could be implemented in different ways, which would impact performance evaluations. In addition, it is common for ML model evaluations to be impaired by circularity and data leakage between the training and test sets [6], which in the original study might have happened at multiple pipeline steps including missing value imputation, feature selection, feature normalization, or hyper-parameter optimization. Such issues can artificially inflate AUC values measurably. Our study reiterates the need to adopt more detailed reporting practices (reporting checklists were published for ML [21] and MRI [22] studies) and to publicly share all software used in the analyses.

In addition to checklists, a possible option to improve reporting practices is to implement analyses using Jupyter notebooks and share them through an online version control platform such as GitHub. We are aware of the pitfalls of Jupyter notebooks regarding good coding practices and potential hidden execution states creating ambiguity. Nevertheless, we believe that their ability to mix code, data, and natural text makes them an excellent platform to report data analyses. We implemented such a notebook for our analysis (available at https://github.com/LivingPark-MRI/shu-etal), which turned out to be more challenging than initially anticipated due to the fact that (1) the PPMI data usage agreement forbid data redistribution and (2) the dataset was not accessible through an application programming interface (API). Therefore, our notebook had to access PPMI by controlling a web browser, which is not maintainable in the long run. To facilitate the development of similar notebooks in the future, we recommend that data sharing initiatives that must prevent data redistribution provide a mechanism to access and query the data programmatically.

Finally, we acknowledge the importance of involving the original authors in reproduction or replication studies. We contacted the senior author of [1] to seek their feedback on an initial draft of this manuscript. Our exchanges confirmed that the discrepancies in classification performance between our models and the one reported in [1] are likely due to (1) undocumented variations in cohort construction, (2) differences in radiomics feature extraction between the A.K software and PyRadiomics, and (3) variations in ML model construction (in particular hyper-parameter optimization) and evaluation, aggravated by the use of a relatively small sample size. This feedback reiterates our previous recommendations on data versioning, sample sizes, reporting practices, and the use of publicly-available software to improve the reproducibility of MRI-based ML studies.

## Acknowledgments

This work was funded by the Michael J. Fox Foundation for Parkinson’s Research (MJFF-021134). We warmly thank Dr. Minming Zhang for providing feedback on the manuscript.

## Notes

### Competing Interest Statement

The authors have declared no competing interest.

